# Influence of Activating and Inhibitory Killer Immunoglobulin-Like Receptors (KIR) genes on the recurrence rate of ocular toxoplasmosis in Brazil

**DOI:** 10.1101/2020.09.16.299446

**Authors:** Daiana de Souza Perce-da-Silva, Thays Euzebio Joaquim, Ana Luisa Quintella do Couto Aleixo, Juliana Motta, Marcelo Ribeiro-Alves, Joseli de Oliveira Ferreira, Luís Cristóvão de Moraes Sobrino Porto, Dalma Maria Banic, Maria Regina Reis Amendoeira

## Abstract

**Background:** Recurrence is a hallmark of ocular toxoplasmosis (OT), and conditions that influence its occurrence remain a challenge. Natural killer cells (NK) are effectors cells whose primary function is the cytotoxic activity against many parasites, including *Toxoplasma gondii*. Among the NK cell receptors, immunoglobulin-like receptors (KIR) deserve attention due to their high polymorphism. This study aimed to analyze the influence of KIR gene polymorphism in the course of OT infection and its association with recurrences after an active episode.

**Methods:** Ninety-six patients from the Ophthalmologic Clinic of the National Institute of Infectology Evandro Chagas (INI/Fiocruz/RJ, Brazil) were followed for up to five years. After DNA extraction, genotyping of the patients was performed by PCR-SSO utilizing Luminex equipment for reading. During follow-up, 57.4% had a recurrence.

**Results:** We identified 25 KIR genotypes and found a higher frequency of genotypes 1 (31.7%) with worldwide distribution. We note that the *KIR2DL2* inhibitor gene and the gene activator *KIR2DS2* were more frequent in patients without recurrence (P = 0.03 and P = 0.02, respectively). Additionally, we observed one activating gene, KIR2DS1, associated with more than four times faster progression to the development of recurrent ocular toxoplasmosis than individuals without this gene (aRR = 4.6, P = 0.04).

**Conclusion:** The KIR2DL2 and KIR2DS2 are associated as possible protection markers and the KIR2DS1 acting as a possible susceptibility marker. Additionally, the lower proportion of activating genes observed in individuals with recurrence corroborating with the hypothesis that these individuals are more susceptible to ocular toxoplasmosis recurrence (OTR).

## Introduction

*Toxoplasma gondii* is an obligate intracellular protozoan parasite that belongs to the phylum apicomplexa, subclass coccidia. The parasite has a worldwide distribution with a high prevalence that infects humans, birds, rodents, and other animals (intermediate hosts) and felids (definitive hosts) on all continents. The toxoplasmosis can be classified as congenital or acquired. Congenital toxoplasmosis may be exclusively ocular or accompanied by systemic or central nervous system changes [1]. In recent decades, advances show that toxoplasmosis is one of the most important causes of posterior uveitis in the world, representing up to 85% of all cases [1–3]. The ocular lesions are characterized by necrotizing retinitis with oval or circular lesions. Besides it, the lesion can remain active for weeks, and even after healing, it may contain *T. gondii* cysts, so the protozoan remains viable in tissues for years [4].

Ocular toxoplasmosis (OT) is a disease characterized by recurrence episodes. However, the conditions associated with recurrence episodes have not been completely elucidated. After infection, the ocular symptoms depend on complex and variable factors, such as socioeconomic factors and the parasite genotype. In Brazil, both the prevalence (> 80%) and severity of OT are higher than in many other parts of the world [2].

Concerning the immune system, in the eye, the parasite contrives to manipulate the immune response in such a way as to favor its survival without causing too much damage to the organ [2, 5]. In the early phase of *T. gondii* infection, innate immunity cells are recruited to the site of infection. Natural Killer (NK) are important lymphocytes acting in the acute phase of the toxoplasmosis [6]. *In vivo* studies revealed that control of *T. gondii* requires the early production of the pro-inflammatory cytokine IL-12, which stimulates NK, CD4^+^, and CD8^+^ lymphocytes to release IFN-γ [6–8].

Control of NK cell action is through membrane receptors, including Killer Immunoglobulin-Like Receptors (KIR), which recognize Human leukocyte antigen class I molecules (HLA class I) expressed by most cells in the body. The extensive genetic polymorphism of KIR receptors and the regulation of their expression in different NK cell clones are essential factors that delineate each individual’s innate and adaptive immune response.

NK cells have great importance in the control of *T. gondii* infection however, the role of *KIR* genes that encode the immune receptors of NK cells and can trigger local inflammation in the eye has not been elucidated in ocular toxoplasmosis yet. *KIR* genes have been described as risk or protective factors in different types of inflammatory ocular diseases [9–11] and are also associated with many other infectious diseases [12–19].

In ocular toxoplasmosis, no study has evaluated the association of KIR receptors with ocular toxoplasmosis involving recurrence events. Thus, the characterization of these receptors in individuals with ocular toxoplasmosis may help to understand their role in regulating the immune response, clinical evolution of the disease, as well as their relationship with faster or lower recurrences. Besides it, the identification of predisposal individuals may help in their clinical management. However, others studies should be performed such as histological analyses of the ocular tissue affected by *T. gondii* and NK cytotoxicity assays to better understanding the role of NK cells and the expression of KIR in the immunopathogenesis of ocular toxoplasmosis.

## Material and methods

### Ethics statement

The Research Ethics Committee of the National Institute of Infectology Evandro Chagas (INI/Fiocruz) has approved this study protocol as a subproject under the register CAAE 0075.0.009.000-11. All the volunteers gave written informed consent after being informed about the study’s nature, including the objectives, laboratory procedures that would be performed. They allowed the store and future use in research of their samples.

### Patients

This study was conducted using 96 blood and serum samples stored at the Toxoplasmosis Laboratory at IOC-Fiocruz. The patients were attended by the same ophthalmologist between January 2010 and January 2014, and follow-up until July 2015 at the outpatient unit of the Infectious Ophthalmology Laboratory of the National Institute of Infectology Evandro Chagas at Fiocruz [20].

For the purpose of this study, the patients were classified according to the recurrence of ocular toxoplasmosis. As previously described in Aleixo *et. al* [20, 21], recurrent cases were defined as an active retinochoroiditis associated with retinal scar in either eye [22]. It’s important to highlight that episodes of inflammation of the anterior segment in eyes with retinochoroiditis scars were not considered recurrence [23]. Creamy-white focal retinochoroidal lesions in the absence of other retinochoroidal scars were considered as primary lesions. Primary retinochoroidal lesion cases with no recurrence were considered to be highly probable of ocular toxoplasmosis and thus included in the follow-up [21]. The follow-up criteria adopted for the inclusion of patients in the non-recurrence group are described in more detail in Aleixo et al. [20]. It has been demonstrated that the risk of OT recurrence is higher in the year following the first infection than in future years [20, 24]. However, to avoid erroneous associations we included in the non-recurrence group only patients who were followed up for at least two years.

The patients’ exclusion criteria’s were pregnant during any recurrent episodes, genetically related, having co-morbidities (e.g., chronic renal failure), systemic infections (e.g., AIDS, syphilis, and tuberculosis), autoimmune diseases, history of intravenous drugs use, single and unilateral or multiple exudative lesions of retinochoroiditis, and history of cancer chemotherapy or immunosuppressive drug or peri- and/or intraocular steroids use. All exams were performed in the Laboratory of Immunology and Immunogenetics (INI/Fiocruz).

### Genomic DNA extraction

According to the manufacturer’s instructions, genetic DNA was isolated from peripheral blood samples collected in EDTA by using a QIAamp^®^ DNA Blood Midi/Maxi Kit (Qiagen, Valencia, CA). The DNA concentration was determined using a Qubit fluorimeter (Life Technologies, Carlsbad, CA), and the filtrates containing the isolated DNA were stored at −20° C until time of use.

### *KIR* genotyping

The reverse sequence-specific oligonucleotide technique (One Lambda Inc., Canoga Park, CA) with Luminex xMap technology (Luminex Corp., Austin, TX) was used for the typing of the 14 KIR genes and two KIR pseudogenes namely *KIR2DL1, KIR2DL2, KIR2DL3, KIR2DL4, KIR2DL5, KIR2DS1, KIR2DS2, KIR2DS3, KIR2DS4, KIR2DS5, KIR3DL1, KIR3DL2, KIR3DL3, KIR3DS1, KIR2DP1,* and KIR3DP1, according to the manufacturer’s instructions. The carrier frequency was calculated through the direct counting of individuals owning at least one copy of the gene. It was necessary to consider that all frequencies were in Hardy-Weinberg equilibrium (HWE). In contrast, individuals heterozygous for a KIR gene could not be distinguished from those that were homozygous without family members. The genotypic frequency (GF) was calculated by the formula GF=1−√(1−CF), where CF indicates the carrier frequency, calculated by CF=frequency(F)%/100 [25]. The survey data were recorded and entered into an Epi Info 2007 (Centers for Disease Control and Prevention, Atlanta, GA) database.

### Statistical analysis

Mann-Whitney U tests were used to compare clinical-demographic features for continuous numerical variables between recurrent and non-recurrent groups. In contrast, for categorical nominal variables, Fisher’s exact tests were used to evaluate frequencies independence among these features and the groups. Proportional tests based on the chi-square distribution with one degree-of-freedom were used to test if proportions of the non-recurrent group were equal to that of the recurrent group. For each participant, person-years (py) at risk were calculated between the discharge of a previous episode and the discharge date of the last episode of ocular toxoplasmosis or the last follow-up visit, which occurred first. Recurrence rates of ocular toxoplasmosis per 100 py and its ninety-five percent confidence intervals (CI95%) were estimated according to asymptotic standard errors calculated from a Gamma distribution [26]. The effects of potential hazard factors on OTR were assessed using relative risks (RR) and its CI95% estimated by the fit of Simple and Multiple (aRR) Poisson Generalized Linear Mixed Effect Models (P-GLMM) [27] optimized by maximum likelihood. Repeated measurements were treated as random effects. Degrees of freedom for estimated effects were approximated by the Satterthwaite method [28]. Confounding variables were selected by Simple/Bivariate P-GLMM and included in multivariate models if any p-value < 0.2 to eliminate sampling bias. All statistical analyses were performed using R version 3.6.1 [29].

## Results

### General features of the studied population

The clinical-demographic data summarized in Table 1 show that patients with ocular toxoplasmosis were followed up for different periods. In our studied population, most individuals were adults, residents of the state of Rio de Janeiro, and they were followed up over two years. The age range was 14–67 years, with an average of 32 years, and the proportion of men was higher (56.2%) than for women (43.8%). The average follow-up time between recurrence group and non-recurrence group was very similar (1338 and 1370 days, respectively).

**Table 1.** Clinical-demographic features of patients consulted at the outpatient unit of the Infectious Ophthalmology Laboratory of the National Institute of Infectology Evandro Chagas, Fiocruz, RJ (N =96)

|  |  |  |
| --- | --- | --- |
| **Gender** | Male (N/%) | 54/56.2 |
|  | Female (N/%) | 42/43.6 |
|  | Total (N) | 96 |
| Age ( $\bar{x} \pm SD$) | | 32 ± 11 |
| **Follow-up period** | $\leq 2$ years (N/%) | 4/4.2 |
| | $> 2$ and $\leq 3$ years (N/%) | 18/18.7 |
| | $> 3$ and $\leq 4$ years (N/%) | 28/29.2 |
| | $> 4$ and $\leq 5$ years (N/%) | 45/46.9 |
| | $> 5$ and $\leq 6$ years (N/%) | 1/1 |
| **N recurrence** | 0 (N/%) | 38/39.6 |
|  | 1 (N/%) | 33/32.7 |
|  | 2 (N/%) | 18/17.8 |
|  | 3 (N/%) | 5/5.2 |
|  | 5 (N/%) | 2/2.1 |
| **Follow-up in days**<br>( $\bar{x} \pm SD$) | Recurrence group | 1338 ± 324 |
| | Non-recurrence group | 1370 ± 321 |
| **State** | Rio de Janeiro (N/%) | 87/90.6 |
|  | Others* (N/%) | 9/9.4 |

$\bar{x}$ mean; N, number of individuals; SD, standard deviation.; \*Alagoas, Espírito Santo, Mato Grosso, Minas Gerais, Paraíba, and Rio Grande do Sul.

The number of recurrences varied between 1 and 5 episodes, and, according to this variation, 33 individuals (34.4%) had one recurrence episode, 18 (18.7%) had two episodes, 5 (5.2%) presented three episodes and 2 (2.1%) had five recurrence episodes during the follow-up (Table 1).

### KIR genes frequencies

The distribution of KIR gene frequencies (*F*%) in patients consulted at the outpatient unit of the Infectious Ophthalmology Laboratory from INI is illustrated in Table 2. All 16 KIR genes investigated (14 genes and two pseudogenes, including framework loci) were detected in the study population, and the framework loci *KIR2DL4, KIR3DL2, KIR3DL3,* and *KIR3DP1* were present in all individuals. In general, the frequencies of inhibitory KIR genes were higher than 90%, except for the *KIR2DL2* and *KIR2DL5* genes (54.2% and 54.5%, respectively). Also, the proportion of patients carrying any activating KIR gene ranged from 18.7% to 51%, except for the KIR2DS4, with a 96.9% frequency.

**Table 2.** Distribution of KIR gene frequencies in patients consulted at the of the Infectious Ophthalmology Clinic of the National Institute of Infectology Evandro Chagas, Fiocruz, RJ.

| *KIR Gene* | *(N)* | *F%* | *GF* |
| --- | --- | --- | --- |
| *Inhibitory* |  |  |  |
| *2DL1* | 94 | 97.9 | 0.856 |
| *2DL2* | 52 | 54.2 | 0.324 |
| *2DL3* | 88 | 91.7 | 0.712 |
| *2DL5* | 53 | 55.2 | 0.331 |
| *3DL1* | 95 | 98.9 | 0.896 |
| *Activating* |  |  |  |
| *2DS1* | 40 | 41.7 | 0.237 |
| *2DS2* | 49 | 51 | 0.3 |
| *2DS3* | 18 | 18.7 | 0.099 |
| *2DS4* | 93 | 96.9 | 0.824 |
| *2DS5* | 41 | 42.7 | 0.244 |
| *3DS1* | 34 | 35.4 | 0.197 |
| *Framework* |  |  |  |
| *2DL4* | 96 | 100 | 1 |
| *3DL2* | 96 | 100 | 1 |
| *3DL3* | 96 | 100 | 1 |
| *<sup>A</sup>3DP1* | 96 | 100 | 1 |
| *Pseudogene* |  |  |  |
| *2DP1* | 93 | 96.9 | 0.824 |

*F%*, KIR genes frequencies; *GF*, genotypic frequency; KIR, killer cell immunoglobulin-like receptors;
*N*, number of individuals; *A*, framework, and pseudogene.

### *KIR* Genotypes

A total of 24 different KIR genotypes were identified in the studied population (Fig 1) based on the presence and absence of 16 KIR genes. All of these were previously described in the KIR genotype database (www.allelefrequencies.net) and contained between 8 and 16 KIR genes per individual. The genotype 1, the most common, accounted for 31.3% of all genotypes in our studied population, and genotypes 4, 2 and 3 were detected in 12.5%, 11.5%, and 8.3% of the studied population respectively (Figure 1). Among the 24 genotypes, more than half (14 genotypes) were in low frequency, presenting only in 1% of the population. The two most common genotypes (1 and 4) observed in our population were present at a similar frequency to that of North, Central, and South American populations (www.allelefrequencies.net) (*P* > 0.05 for all). The population was classified according to the haplogroups (A and B) existing for the KIR genes. Haplogroup B was named Bx because of the lack of distinction between the haplotype AB and BB. The haplogroup Bx was the most frequent (68.7%), while the haplogroup AA was present in 31.3% of our population.

**Figure 1.**
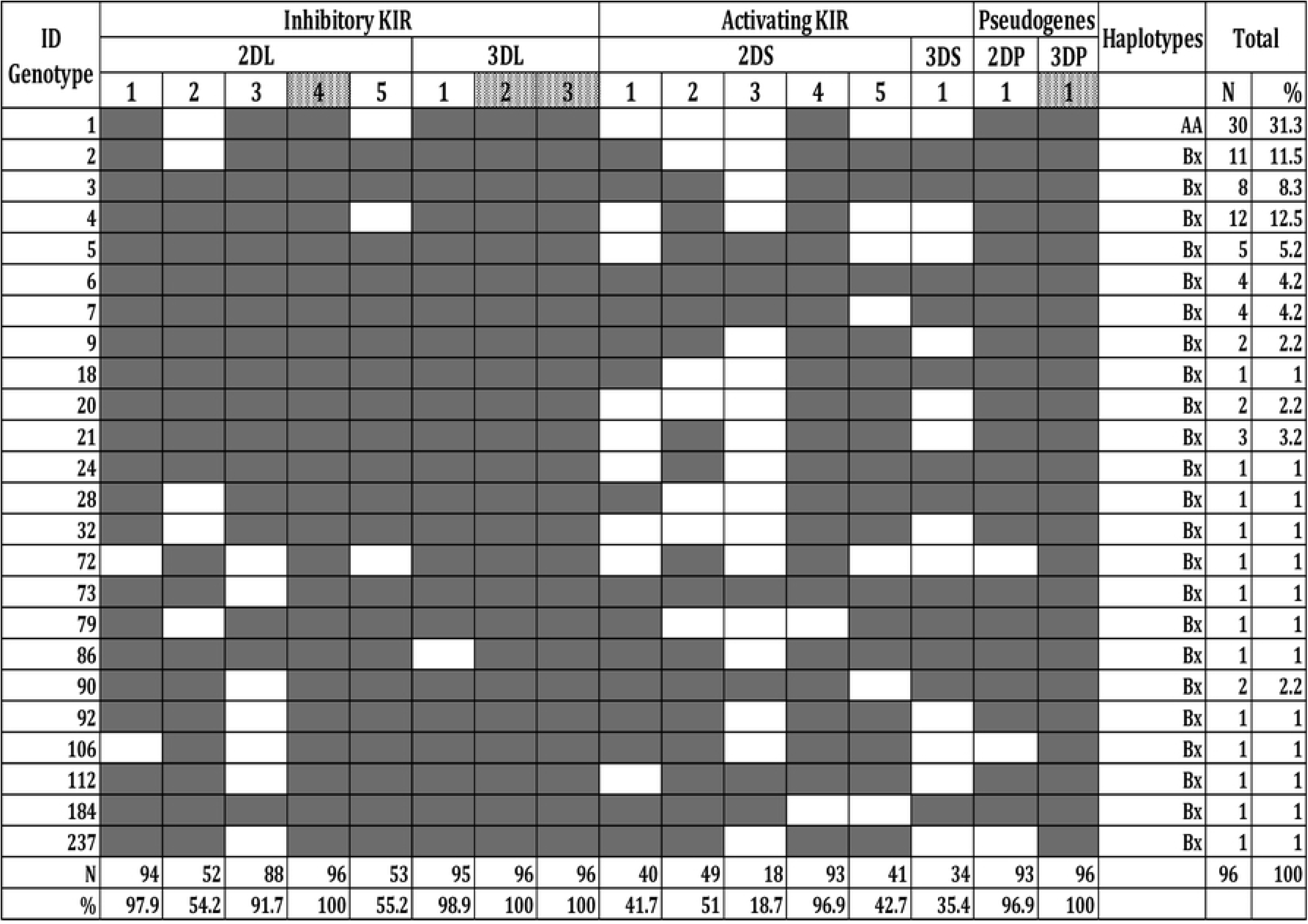

### Comparison of KIR genes frequencies between the recurrence and non-recurrence group

In order to evaluate the possible influence of KIR genes on the recurrence of ocular toxoplasmosis after active episode, the recurrence group (58/60.4%) was compared to the non-recurrence group (38/39.6%). About the follow-up time, more than half (32/55%) of the individuals in the recurrence group were followed for more than four years. In the non-recurrence group, the majority (26/68.5%) of the individuals were followed for more than three years. When we evaluated if the difference between KIR gene frequencies among groups were significant, we observed that between the 16 genes KIR evaluated, the inhibitory *KIR2DL2* and the activating *KIR2DS2* were more frequent in the non-recurrence group than in the recurrence group (P = 0.03 and P = 0.02, respectively) (Table 3).

**Table 3.** Distribution of KIR gene frequencies among recurrence and non-recurrence group from an outpatient unit of the Infectious Ophthalmology Laboratory of the National Institute of Infectology Evandro Chagas, Fiocruz, RJ.

| KIR Receptor | Recurrence group<br>(N= 58)<br>N/% | Non-recurrence<br>group (N= 38)<br>N/% | P-Value |
| --- | --- | --- | --- |
| ***Inhibitory*** |  |  |  |
| *KIR2DL1* | 57/98.3 | 37/97.3 | 1 |
| *KIR2DL2* | 26/45 | **26/68.4** | **0.03** |
| *KIR 2DL3* | 53/91.4 | 35/92.1 | 1 |
| *KIR 2DL5* | 33/57 | 20/53 | 0,8 |
| *KIR 3DL1* | 58/100 | 37/97.3 | 0,3 |
| ***Activating*** |  |  |  |
| *KIR2DS1* | 26/45 | 14/37 | 0,5 |
| *KIR2DS2* | 24/41.4 | **25/66** | **0.02** |
| *KIR2DS3* | 18/31 | 8/21 | 0.3 |
| *KIR2DS4* | 57/ 98.3 | 36/95 | 0.5 |
| *KIR2DS5* | 26/45 | 15/39 | 0.6 |
| *KIR3DS1* | 21/36.2 | 13/34.2 | 1 |
| ***Framework*** |  |  |  |
| *KIR2DL4* | 58/100 | 38/100 | - |
| *KIR3DL2* | 58/100 | 38/100 | - |
| *KIR3DL3* | 58/100 | 38/100 | - |
| *<sup>a</sup>KIR3DP1* | 58/100 | 38/100 | - |
| ***Pseudogene*** |  |  |  |
| *KIR2DP1* | 56/96.5 | 37/97.4 | 1 |

N, number of individuals; %, KIR genes frequencies; <sup>a</sup>, framework, and pseudogene.

When we evaluated a possible association between KIR genes and the acceleration of recurrence of ocular toxoplasmosis after an active episode, we used the multiple analysis to estimate the relative-risks of recurrence. We observed that recurrence was 4.6 times in individuals with KIR2DS1 compared with individuals without this gene (Table 4). Besides, it is interesting to mention that recurrence was 1.2 faster in individuals between 20 and 40 years old.

**Table 4.** Poisson Generalized Linear Mixed Effect Models (P-GLMM) for adjusted incidence ratios between KIR gene carriers/non carriers and the time until the recurrence of ocular toxoplasmosis.

| Features | Levels | Outcome | pY | Incidence/100 pY<br>(CI95%) | aRR (CI95%);<br>p-value |
| --- | --- | --- | --- | --- | --- |
| Overall |  | 94 | 359,2087611 | 26.17 (21.15-32.02) |  |
| Gender | M | 58 | 196,36 | 29.54 (22.43-38.18) | Reference |
|  | F | 36 | 162,85 | 22.11 (15.48-30.6) | 2.145 (1.6526-3.1879); 0.2052 |
| Age | (20,40] | 69 | 252,87 | 27.29 (21.23-34.53) | Reference |
|  | (0,20] | 2 | 34,67 | 5.77 (0.7-20.84) | **1.2534 (1.0565-2.5306); 0.0391** |
|  | (40,67] | 23 | 71,68 | 32.09 (20.34-48.15) | 3.0026 (1.9814-5.8579); 0.6956 |
| KIR2DL1 | 1 | 93 | 351,98 | 26.42 (21.33-32.37) | Reference |
|  | 0 | 1 | 7,23 | 13.84 (0.35-77.11) | 2.8038 (1.1485-2161.2466); 0.9762 |
| KIR2DL2 | 1 | 39 | 195,39 | 19.96 (14.19-27.29) | Reference |
|  | 0 | 55 | 163,82 | 33.57 (25.29-43.7) | 2.4239 (1.3767-11.6106); 0.8148 |
| KIR2DL3 | 1 | 89 | 334,18 | 26.63 (21.39-32.77) | Reference |
|  | 0 | 5 | 25,03 | 19.97 (6.49-46.61) | 2.6406 (1.4562-12.2912); 0.9516 |
| KIR2DL5 | 1 | 56 | 202,75 | 27.62 (20.86-35.87) | Reference |
|  | 0 | 38 | 156,46 | 24.29 (17.19-33.34) | 1.9611 (1.5333-2.8899); 0.0884 |
| KIR2DP1 | 1 | 92 | 347,51 | 26.47 (21.34-32.47) | Reference |
|  | 0 | 2 | 11,7 | 17.09 (2.07-61.74) | 2.5209 (1.245-49.4396); 0.915 |
| KIR2DS1 | 0 | 49 | 209,58 | 23.38 (17.3-30.91) | Reference |
|  | 1 | 45 | 149,63 | 30.07 (21.94-40.24) | 4.6253 (2.7393-10.2543); 0.0459 |
| KIR2DS2 | 1 | 35 | 183,39 | 19.08 (13.29-26.54) | Reference |
|  | 0 | 59 | 175,81 | 33.56 (25.55-43.29) | 5.8398 (1.8667-146.8155); 0.2841 |
| KIR2DS3 | 0 | 80 | 295,18 | 27.1 (21.49-33.73) | Reference |
|  | 1 | 14 | 64,03 | 21.87 (11.95-36.69) | 3.209 (1.8048-9.9995); 0.6583 |
| KIR2DS4 | 1 | 93 | 347,99 | 26.72 (21.57-32.74) | Reference |
|  | 0 | 1 | 11,21 | 8.92 (0.23-49.68) | 1.5681 (1.0642-25.8153); 0.4286 |
| KIR2DS5 | 0 | 48 | 203,24 | 23.62 (17.41-31.31) | Reference |
|  | 1 | 46 | 155,97 | 29.49 (21.59-39.34) | 3.854 (2.4221-7.8259); 0.1643 |
| KIR3DS1 | 0 | 56 | 233,58 | 23.97 (18.11-31.13) | Reference |
|  | 1 | 38 | 125,63 | 30.25 (21.4-41.52) | 3.9636 (2.4694-8.1504); 0.1363 |

### Analysis with the weighted average of Inhibitory and Activating KIR between recurrence and non-recurrence group

In order to evaluate if the weighted mean of inhibitory and activating genes present in the genotypes found in our population would be associated with recurrence of ocular toxoplasmosis, we used the formula WM = [(ATIV * 5) - (INIB * 3)], where WM is the weighted mean of inhibitory and activating genes, ATIV is the number of activating KIR genes, and INIB is the number of inhibitory KIR genes of an individual. With this calculation, we verified that the negative weighted mean belonged to individuals with a lower proportion of activating genes (between 1 and 4 genes, N = 75). In comparison, the positive weighted mean was present in individuals with a higher proportion of activating genes (between 4 to 6 genes, N = 21).

We observed in our population that the majority of individuals had a lower number of activating KIR genes (1 to 4 activator genes) (78%; P < 0.01). Among these individuals, the majority belonged to the group of individuals with recurrence of ocular toxoplasmosis (62.7%, P = 0.003) (Table 5).

**Table 5.** The weighted mean of inhibitory and activating KIR between recurrence and non-recurrence groups from an outpatient unit of the Infectious Ophthalmology Laboratory of the National Institute of Infectology Evandro Chagas (INI), Fiocruz, RJ.

| Weighted mean | Individuals |  | P-Value |
| --- | --- | --- | --- |
|  | Recurrence | Non-recurrence |  |
|  | group | group |  |
|  | (N/%) | (N/%) |  |
| Negative (1 to 4 activating KIR) (N= 75) | 47/62.7 | 28/37.3 | **0.003** |
| Positive (4 to 6 inhibitory KIR) (N=21) | 11/52.4 | 10/47.6 | 1 |

*N*, number of individuals; %, KIR genes frequencies;

## Discussion

Because of KIR’s role in the immune response and its extensive genomic diversity, it is known that variation in KIR genes affects the resistance and/or susceptibility to the pathogenesis of various diseases. In this context, the current study characterized the gene frequency profile of the 16 genes encoding KIR receptors and evaluated their influence in patients with recurrence and non-recurrence ocular toxoplasmosis after active episodes.

In our study, 91% of the 101 participants were followed for at least two years. The number of recurrences varied from 1 to 5 episodes. About the follow-up time, more than half (55%) of the individuals in the recurrence group were followed for more than four years. In the non-recurrence group, the majority (68.5%) of the individuals were followed for more than three years.

Concerning the profile of the gene frequencies of KIR receptors in the study population, we observed that the genes KIR3DL1, KIR2DL1, KIR2DS4, and KIR2DP1 were the most prevalent (above 97%). These high frequencies are similar to those found in other Brazilian populations, including Rio de Janeiro [19, 25, 30–33].

When we evaluated whether there were significant differences in the frequencies of the KIR genes between the recurrence and non-recurrence group, we observed that the inhibitory gene KIR2DL2 and the activator gene KIR2DS2 were more frequent in the non-recurrence group than in recurrence group (P = 0.03 and P = 0.02, respectively). According to the literature, the KIR2DL2 and KIR2DS2 genes are in strong linkage disequilibrium in most populations worldwide [25, 34, 35]. Due to this strong linkage disequilibrium between the two loci, it is difficult to determine which one mediates the effect [36].

The inhibitor gene KIR2DL2 and its corresponding activator KIR2DS2 have already been associated with other diseases. They have been associated with protection against HIV-1 infection [37], and KIR2DL2+HLA-C1 was associated with a decreased risk of chronic HBV [13]. In contrast, a previous study that determined and compared the frequencies of the KIR genes of children with severe or uncomplicated malaria with healthy controls in the same area found that the frequencies of both genes were significantly higher in malaria cases (severe or uncomplicated) than in controls. These results suggest that KIR2DL2 + C1 and/or KIR2DS2 + C1 carriers were more at risk of being infected with malaria parasites than those without any of these genotypes [36]. As in our population, both genes were more frequent in the non-recurrence group. We can suggest that the inhibitory gene KIR2DL2, together with the activator KIR2DS2, may be acting as protective markers, slowing the recurrence of ocular toxoplasmosis.

Regarding the time until the recurrence of ocular toxoplasmosis, we found that the KIR2DS1 activating gene accelerated recurrences of ocular toxoplasmosis. Previous studies have shown the involvement of the KIR2DS1 activating gene in inflammatory bowel disease, like Crohn’s disease and ulcerative colitis [38, 39]. On the other hand, Ayo et al. [40] demonstrated an association between others activating KIR and their HLA ligands (KIR3DS1-Bw4-80Ile and KIR2DS1+/C2++ KIR3DS1+/Bw4-80Ile+) and increased susceptibility for ocular toxoplasmosis and its clinical manifestations.

NK cells can express from one to all possible KIR genes present in their genotype on their surfaces. Considering that the KIR repertoire depends directly on the inherited KIR genes, the genotype and allelic diversities of KIR genes can affect the function and the levels of specificity and sensitivity of NK cells [41]. In this context, we evaluated whether the variation in the proportion between inhibitory and activating KIR genes, present in the genotypes of our population, could be associated with OTR.

In the analysis of the frequencies of the activating genes in the studied population, we observed that the majority of individuals (78%) have a lower proportion of activating KIR genes (between 1 to 4 activating genes) (P < 0.01). As for the distribution of individuals who presented a lower proportion of activating genes, we observed that the majority belonged to individuals with recurrence (62.7%; P = 0.003). This finding is in agreement with the foundation for the balance of the immune response. The balance between the inhibition and activation signals regulate NK cells and is responsible for protecting or predisposing an individual to diseases [42–44]. It allows us to suggest that the lower proportion of activating KIR genes is associated with accelerating OTR episodes.

In the present study, we characterize KIR genes’ profile in patients with ocular toxoplasmosis after an active episode. We observed two genes (KIR2DL2 and KIR2DS2), acting together as possible protection markers. Additionally, we founded one activating gene (KIR2DS1) acting as a possible susceptibility marker and the lower proportion of activating genes observed in individuals with recurrence corroborating with the hypothesis that these individuals are more susceptible to OTR. As far as we know, no literature relates to the polymorphism of KIR and the follow up of recurrence of ocular toxoplasmosis for a prolonged period (up to 5 years). Thus, the characterization of KIR genes makes this study a pioneer in searching for an association between the KIR genes polymorphisms and the time until recurrence of ocular toxoplasmosis.

## Author Contributions Conceptualization

**Conceived and designed the experiments:** DSPS, MRRA, DMB, JOF, LCMS

**Performed the experiments:** DSPS, TEJ, ALQC, JM, LCMSP

**Analyzed the data:** DSPS, TEJ, MRA, JOF, DMB, LCMSP, MRRA

**Contributed reagents/materials/analysis:** MRRA, ALQC, LCMS

**Wrote the paper:** DSPS, TEJ, DMB, MRRA

## References

1. Maenz M, Schlüter D, Liesenfeld O, Schares G, Gross U, Pleyer U. Ocular toxoplasmosis past, present and new aspects of an old disease. Prog Retin Eye Res. 2014 Mar;39:77–106.

2. Garweg JG. Ocular Toxoplasmosis: an Update. Klin Monbl Augenheilkd. 2016 Apr;233(4):534–9.

3. Talabani H, Mergey T, Yera H, Delair E, Brézin AP, Langsley G, et al. Factors of occurrence of ocular toxoplasmosis. A review. Parasite. 2010 Sep;17(3):177–82.

4. Smith JR, Ashander LM, Arruda SL, Cordeiro CA, Lie S, Rochet E, Belfort R, Furtado JM. Pathogenesis of ocular toxoplasmosis. Prog Retin Eye Res. 2020 Jul 24:100882.

5. Sultana MA, Du A, Carow B, Angbjär CM, Weidner JM, Kanatani S, et al. Down modulation of Effector Functions in NK Cells upon *Toxoplasma gondii* Infection. Infect Immun. 2017 Sep 20;85(10):e00069–17

6. Sasai M, Yamamoto M. Innate, adaptive, and cell-autonomous immunity against *Toxoplasma gondii* infection. Exp Mol Med. 2019 Dec 11;51(12):1–10.

7. Khan IA, Matsuura T, Kasper LH. Interleukin-12 enhances murine survival against acute toxoplasmosis. Infect Immun. 1994 May;62(5):1639–42.

8. Gazzinelli RT, Hieny S, Wynn TA, Wolf S, Sher A. Interleukin 12 is required for the T-lymphocyte-independent induction of interferon gamma by an intracellular parasite and induces resistance in T-cell-deficient hosts. Proc Natl Acad Sci U S A. 1993 Jul 1;90(13):6115–9.

9. Kim SJ, Lee S, Park C, Seo JS, Kim JI, Yu HG. Targeted resequencing of candidate genes reveals novel variants associated with severe Behçet’s uveitis. Exp Mol Med. 2013;45(10):e49.

10. Levinson RD, Martin TM, Luo L, Ashouri E, Rosenbaum JT, Smith JR, et al. Killer cell immunoglobulin-like receptors in HLA-B27-associated acute anterior uveitis, with and without axial spondyloarthropathy. Invest Ophthalmol Vis Sci. 2010 Mar;51(3):1505–10.

11. Levinson RD. Killer immunoglobulin-like receptor genes in uveitis. Ocul Immunol Inflamm. 2011 Jun;19(3):192–201.

12. Chaisri S, Jumnainsong A, Romphruk A, Leelayuwat C. The effect of KIR and HLA polymorphisms on dengue infection and disease severity in northeastern Thais. Med Microbiol Immunol. 2020 Jun 10.

13. Auer ED, Tong HV, Amorim LM, Malheiros D, Hoan NX, Issler HC, et al. Natural killer cell receptor variants and chronic hepatitis B virus infection in the Vietnamese population. Int J Infect Dis. 2020 Jul;96:541–547.

14. Tuttolomondo A, Di Raimondo D, Vasto S, Casuccio A, Colomba C, Norrito RL, et al. Protective and causative killer Ig-like receptor (KIR) and metalloproteinase genetic patterns associated with Herpes simplex virus 1 (HSV-1) encephalitis occurrence. J Neuroimmunol. 2020 Jul 15;344:577241.

15. Torimiro J, Yengo CK, Bimela JS, Tiedeu AB, Lebon PA, Sake CS, et al. Killer Cell Immunoglobulin-Like Receptor Genotypes and Haplotypes Contribute to Susceptibility to Hepatitis B Virus and Hepatitis C Virus Infection in Cameroon. OMICS. 2020 Feb;24(2):110–115.

16. de Sá NBR, Ribeiro-Alves M, da Silva TP, Pilotto JH, Rolla VC, Giacoia-Gripp CBW, et al. Clinical and genetic markers associated with tuberculosis, HIV-1 infection, and TB/HIV-immune reconstitution inflammatory syndrome outcomes. BMC Infect Dis. 2020 Jan 20;20(1):59.

17. Alves HV, de Moraes AG, Pepineli AC, Tiyo BT, de Lima Neto QA, Santos TDS, et al. The impact of KIR/HLA genes on the risk of developing multibacillary leprosy. PLoS Negl Trop Dis. 2019 Sep 16;13(9):e0007696.

18. Prakash S, Ranjan P, Ghoshal U, Agrawal S. KIR-like activating natural killer cell receptors and their association with complicated malaria in north India. Acta Trop. 2018 Feb;178:55–60.

19. Perce-da-Silva DS, Silva LA, Lima-Junior JC, Cardoso-Oliveira J, Ribeiro-Alves M, Santos F, et al. Killer cell immunoglobulin-like receptor (KIR) gene diversity in a population naturally exposed to malaria in Porto Velho, Northern Brazil. Tissue Antigens. 2015 Mar;85(3):190–9.

20. Aleixo AL, Curi AL, Benchimol EI, Amendoeira MR. Toxoplasmic Retinochoroiditis: Clinical Characteristics and Visual Outcome in a Prospective Study. PLoS Negl Trop Dis. 2016 May 2;10(5):e0004685.

21. Aleixo ALQDC, Vasconcelos C de Oliveira R, Cavalcanti Albuquerque M, Biancardi AL, Land Curi AL, Israel Benchimol E, et al. Toxoplasmic retinochoroiditis: The influence of age, number of retinochoroidal lesions and genetic polymorphism for IFN-γ +874 T/A as risk factors for recurrence in a survival analysis. PLoS One. 2019 Feb 12;14(2):e0211627.

22. Holland GN. Ocular toxoplasmosis: a global reassessment. Part II: disease manifestations and management. Am J Ophthalmol. 2004 Jan;137(1):1–17.

23. Holland GOCG, Belfort R Jr, Remington JS. Toxoplasmosis. In: Pepose JS, Holland GN, Wilhelmus KR, editors. Ocular Infection & Immunity. St Louis, Missouri: Mosby-Year Book; 1996. p.1183–1223.

24. Holland GN. Ocular toxoplasmosis: a global reassessment. Part I: epidemiology and course of disease. Am J Ophthalmol 2003;136(6):973–88. pmid:14644206

25. Single RM, Martin MP, Gao X, Meyer D, Yeager M, Kidd JR, Kidd KK, Carrington M. Global diversity and evidence for coevolution of KIR and HLA. Nat Genet. 2007 Sep;39(9):1114–9.

26. Lehmann EL, Casella G. Unbiasedness. Theory of Point Estimation. 1998; 83–146.

27. Bates D, Mächler M, Bolker B, Walker S. Fitting linear mixed-effects models using lme4. arXiv preprint arXiv. 2014;1406.5823.

28. Satterthwaite FE. An approximate distribution of estimates of variance components. Biometrics. 1946; 2(6):110–4.

29. Team R Core. R: A language and environment for statistical computing. 2013.

30. Pedroza LS, Sauma MF, Vasconcelos JM, Takeshita LY, Ribeiro-Rodrigues EM, Sastre D, et al. Systemic lupus erythematosus: association with KIR and SLC11A1 polymorphisms, ethnic predisposition and influence in clinical manifestations at onset revealed by ancestry genetic markers in an urban Brazilian population. Lupus. 2011 Mar;20(3):265–73.

31. Augusto DG, Zehnder-Alves L, Pincerati MR, Martin MP, Carrington M, Petzl-Erler ML. Diversity of the KIR gene cluster in an urban Brazilian population. Immunogenetics. 2012 Feb;64(2):143–52.

32. Augusto DG, Lobo-Alves SC, Melo MF, Pereira NF, Petzl-Erler ML. Activating KIR and HLA Bw4 ligands are associated to decreased susceptibility to pemphigus foliaceus, an autoimmune blistering skin disease. PLoS One. 2012;7(7):e39991.

33. Franceschi DS, Mazini PS, Rudnick CC, Sell AM, Tsuneto LT, de Melo FC,et al. Association between killer-cell immunoglobulin-like receptor genotypes and leprosy in Brazil. Tissue Antigens. 2008 Nov;72(5):478–82.

34. Denis L, Sivula J, Gourraud PA, Kerdudou N, Chout R, Ricard C, et al. Genetic diversity of KIR natural killer cell markers in populations from France, Guadeloupe, Finland, Senegal and Réunion. Tissue Antigens. 2005 Oct;66(4):267–76.

35. Yindom LM, Leligdowicz A, Martin MP, Gao X, Qi Y, Zaman SM, et al. Influence of HLA class I and HLA-KIR compound genotypes on HIV-2 infection and markers of disease progression in a Manjako community in West Africa. J Virol. 2010 Aug;84(16):8202–8.

36. Yindom LM, Forbes R, Aka P, Janha O, Jeffries D, Jallow M, et al. Killer-cell immunoglobulin-like receptors and malaria caused by Plasmodium falciparum in The Gambia. Tissue Antigens. 2012 Feb;79(2):104–13.

37. Sorgho PA, Djigma FW, Martinson JJ, Yonli AT, Nagalo BM, Compaore TR, et al. Role of Killer cell immunoglobulin-like receptors (KIR) genes in stages of HIV-1 infection among patients from Burkina Faso. Biomol Concepts. 2019 Dec 19;10(1):226–236.

38. Samarani S, Mack DR, Bernstein CN, Iannello A, Debbeche O, Jantchou P, Faure C, Deslandres C, Amre DK, Ahmad A. Activating Killer-cell Immunoglobulin-like Receptor genes confer risk for Crohn’s disease in children and adults of the Western European descent: Findings based on case-control studies. PLoS One. 2019 Jun 13;14(6):e0217767.

39. Fathollahi A, Aslani S, Mostafaei S, Rezaei N, Mahmoudi M. The role of killer-cell immunoglobulin-like receptor (KIR) genes in susceptibility to inflammatory bowel disease: systematic review and meta-analysis. Inflamm Res. 2018 Sep;67(9):727–736.

40. Ayo CM, Frederico FB, Siqueira RC, Brandão de Mattos CC, Previato M, Barbosa AP, et al. Ocular toxoplasmosis: susceptibility in respect to the genes encoding the KIR receptors and their HLA class I ligands. Sci Rep. 2016 Nov 9;6:36632.

41. Rempel JD, Hawkins K, Lande E, Nickerson P. The potential influence of KIR cluster profiles on disease patterns of Canadian Aboriginals and other indigenous peoples of the Americas. Eur J Hum Genet. 2011 Dec;19(12):1276–80.

42. Almeida-Oliveira A, Smith-Carvalho M, Porto LC, Cardoso-Oliveira J, Ribeiro Ados S, Falcão RR, et al. Age-related changes in natural killer cell receptors from childhood through old age. Hum Immunol. 2011 Apr;72(4):319–29.

43. Hoteit R, Abboud M, Bazarbachi A, Salem Z, Shammaa D, Zaatari G, et al. KIR genotype distribution among Lebanese patients with Hodgkin’s lymphoma. Meta Gene. 2015 Mar 25;4:57–63.

44. Hoteit R, Bazarbachi A, Antar A, Salem Z, Shammaa D, Mahfouz R. KIR genotype distribution among patients with multiple myeloma: Higher prevalence of KIR 2DS4 and KIR 2DS5 genes. Meta Gene. 2014 Oct 9;2:730–6.

